# Network-based Machine Learning Approach for Structural Domain Identification in Proteins

**DOI:** 10.1101/2020.02.22.960666

**Authors:** Anirudh Tiwari, Nita Parekh

## Abstract

In the era of structural genomics, with a large number of protein structures becoming available, identification of domains is an important problem in protein function analysis as it forms the first step in protein classification. Domain identification has been an active area of research for over four decades and a wide range of automated methods have been proposed. In the proposed network-based machine learning approach, NML-DIP, a combination of supervised (SVM) and unsupervised (k-means) machine learning techniques are used for domain identification in proteins. The algorithm proceeds by first representing protein structure as a protein contact network and using topological properties, *viz*., length, density, and interaction strength (that assesses inter- and intra-domain interactions) as feature vectors in the first SVM to distinguish between single and multi-domain proteins. A second SVM is used to identify the number of domains in multi-domain proteins. Thus, it does not require a prior information of the number of domains. The *k*-means algorithm is then used to identify the domain boundaries that are assessed using CATH annotation. The performance of the proposed algorithm is evaluated on four benchmark datasets and compared with four state-of-the-art domain identification methods. The performance of the approach is comparable to other domain identification tools and works well even when the domains are formed with non-contiguous segments. The performance of the program is significantly improved for prior information about the number of domains. The algorithm is available at: https://bit.ly/NML-DIP.

## Introduction

Proteins are comprised of domains, folds and motifs, which form its basic building blocks. Genetic recombinant techniques allow reorganization of domains, called domain shuffling, resulting in different combinations of domains in different proteins. This along with swapping and insertion of domains result in complex architectures that are responsible for new protein functions in evolution. Hence, efforts to understand protein evolution and its function have mainly focused on domains as these fold into a stable, semi-independent three-dimensional structure and perform a unique function conserved over the evolution. Protein domains are also very useful in analyzing the mechanisms of protein folding and their stability and structural transformations in various conditions. Being the basic units of protein folding, function and evolution, identification and analysis of domains is the first step in understanding functional and structural aspects of proteins. Although the boundaries of a domain can be determined by visual inspection, there definitely exists a need for developing accurate methods for automatic domain identification as the number of solved protein structures is increasing rapidly. The problem of dividing a protein structure into domains is not yet solved efficiently due to the lack of an unambiguous definition of a domain. The most common definition of domains, based on structural aspects is that these are compact stable modules containing a hydrophobic core and can fold independent of the rest of the protein while the evolutionary and functional aspects of the definition suggest that these can occur in different combinations and perform a specific function (Veretnik *et al* 2009).

Deviations are observed in a number of proteins, such as a domain may be very small and may not contain a hydrophobic core, may occur as a large single structural unit, or two domains together may perform a specific function, instead of each one having its own unique function. This makes computational assignment of domains a difficult task. Existence of non-contiguous domains (occurring as a result of domain insertion) further adds to the difficulty in developing an automated solution for domain partitioning.

Though several methods have been proposed to predict domains they all have notable limitations. Some methods fail to correctly partition non-contiguous domains (Sistla *et al* 2005) or are unable to distinguish between single and multi-domain proteins, while some require specifying the number of domains the protein must be split into. Most domain databases use more than one approach along with manual inspection for correct assignment of domains, for e.g., CATH (Orengo *et al* 1997) uses four domain assignment methods, namely, DETECTIVE (Swindells 1995), DOMAK (Siddiqui and Barton 1995), PUU (Holm and Sander 1994) and the method proposed by Islam *et al* (1995). If all the four methods are unanimous in their identification of domains in a protein, only then the domains are automatically assigned, else a manual judgment is made about the best definition among the four. Structural classification of proteins (SCOP) is another exhaustive database in which the domains are manually (Hubbard *et al* 1999) assigned based on evolutionary and functional relationship. While classification of domains in CATH is based on structural integrity, SCOP focuses on functional and evolutionary aspects, and as a result the two most widely used protein domain databases differ in about 20% of domain assignments. With over 60% of proteins being single domain, and about 25% of multi-domain proteins being non-contiguous, there exists a need for reliable domain identification methods that can handle simple as well as complex domain architectures. An excellent review by Veretnik *et al* (2009) provides comparison of various methods for domain assignment.

## Methodology

The flowchart depicting various steps of the proposed <u>N</u>etwork-based <u>M</u>achine <u>L</u>earning algorithm for <u>D</u>omain Identification in <u>P</u>roteins (NML-DIP) is shown in Figure 1. Here, protein structure is modeled as a Protein Contact Network (PCN), wherein each amino acid residue acts as a node and an edge is drawn between two residues if they are within a distance of 7Å (Di Paola *et al* 2013, Chakrabarty and Parekh 2016). Next, we employ a combination of supervised (SVM) (Hearst 1998) and unsupervised (k-means) (Lloyd 1982) machine learning approaches using graph properties as feature vectors for the SVMs. Various steps in the algorithm are briefly discussed below.

**Fig. 1:**
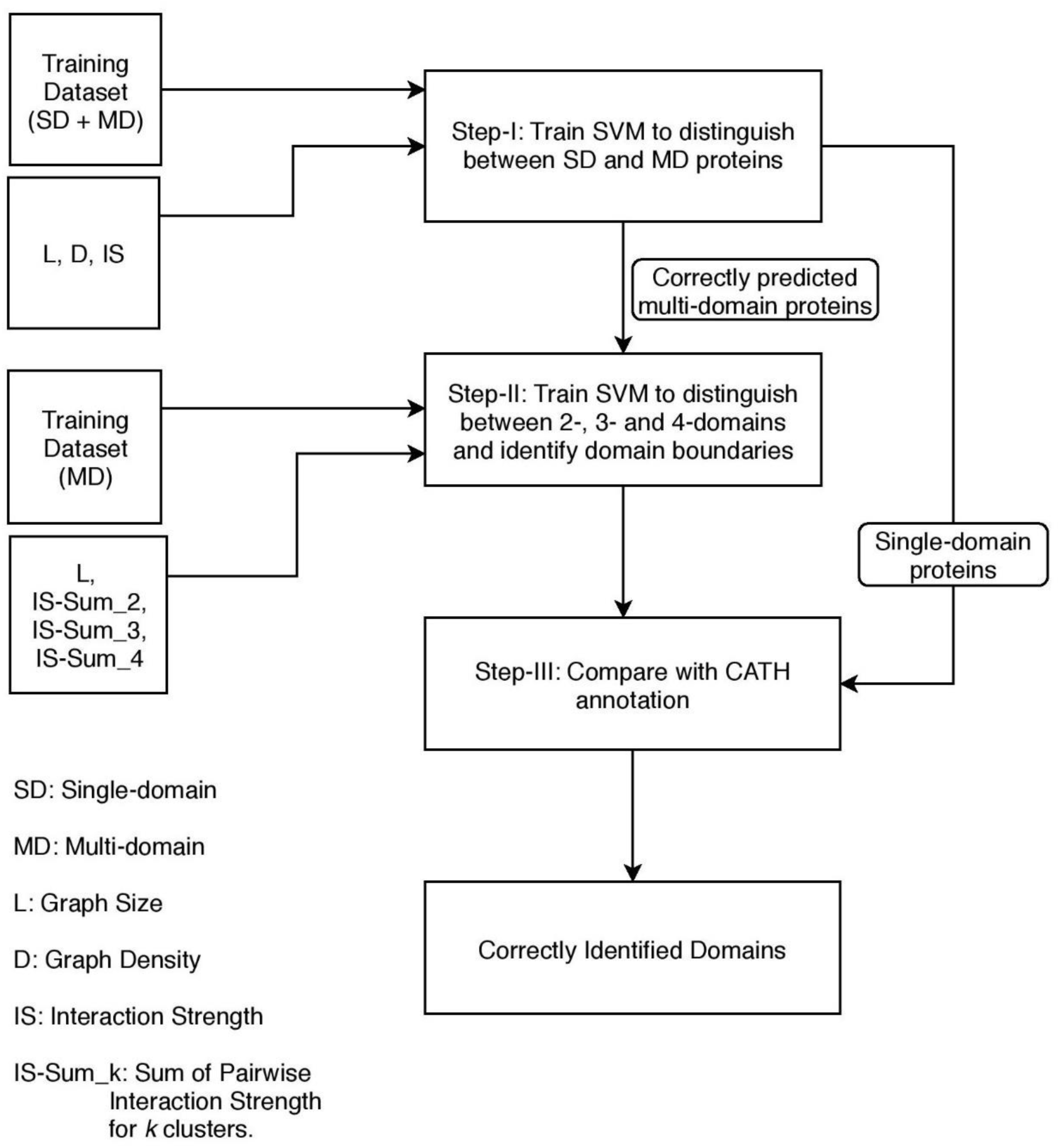
The proposed algorithm, NML-DIP is a 2-step process: first step involves classifying a protein chain as single or multi-domain using SVM. All protein chains correctly classified as multi-domain (based on CATH annotation) are submitted to a second SVM for classifying them as 2-domain, 3-domain, or 4-domain. This step involves k-means for obtaining clusters of size k. In the final step, domain boundaries are obtained using k-means and compared with CATH annotation.

### Step-1: Distinguishing single *vs* multi-domain proteins

The first step in domain classification problem is to identify whether a protein is a single or multi-domain protein. For this task a support vector machine (SVM) is trained using network properties of protein contact network (PCN) as feature vectors, defined below.

### Length of Protein

It is defined as the number of nodes in a PCN (i.e., number of amino acid residues in the protein). The underlying assumption for considering length as one of the features is that the length of protein is expected to increase with increase in the number of domains, though there are notable exceptions.

### Graph Density

It is defined as the ratio of the number of edges, E, observed to the number of possible edges, ½ V(V-1), in a graph of size V, given by Equation 1. Since domains are compact globular structures, graph density is a good measure to quantify the compactness of a protein as a single domain protein is expected to be more compact compared to a multi-domain protein.

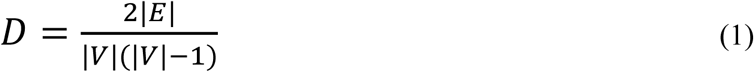

### Interaction Strength

It is well known that the number of inter-domain interactions are fewer compared to intra-domain interactions (Wetlaufer 1973). Thus, if a single-domain protein is split into two domains, we would observe more inter-domain interactions compared to that expected in a natural two-domain protein. These inter-domain interactions are captured by a measure called the interaction strength (IS) (Yalamanchili and Parekh 2009), by computing the number of inter- and intra-domain interactions obtained on splitting a given protein chain into two clusters by k-means algorithm. Mathematically, the Interaction Strength (IS) is defined as:

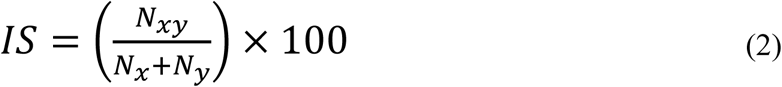

where *N*_*xy*_ are the number of inter-cluster interactions and *N*_*x*_ and *N*_*y*_ are the number of intra-cluster interactions in clusters *x* and *y*, respectively. Here two nodes are said to be interacting if they are within a distance of 7Å and represented by an edge in the PCN.

### Step-2: Identifying Number of Domains, *k*

In this step, a second SVM is used for classifying multi-domain proteins into 2-, 3- or 4-domains. Using k-means, protein chain is split into *k*-clusters (*k* = 2 - 4), i.e., 2-, 3- and 4-domains, and interaction strength (IS) is computed for every cluster-pair in each *k* split. The sum of pairwise Interaction Strengths, IS-Sum_k, between clusters is computed for *k*-split, *k* = 2, 3 and 4, and is expected to increase with increasing *k*. Thus, for *k*=2, IS-Sum_2 = IS-D_1_D_2_, for *k*=3, IS-Sum_3 = IS-D_1_D_2_+IS-D_1_D_3_+IS-D_2_D_3_, and for *k*=4, IS-Sum_4 = IS-D_1_D_2_+IS-D_1_D_3_+IS-D_1_D_4_+IS-D_2_D_3_+IS-D_2_D_4_+IS-D_3_D_4_, where IS-D_i_D_j_ denotes the interaction strength between domains *i* and *j*. If a *k*-domain protein is split into *n* clusters, where *n* > *k*, then IS-Sum_n is expected to be significantly higher than IS-Sum_k as this would lead to one of the true domains getting split and result in higher inter-domain interactions. To capture this information, the second SVM is trained on the IS-Sum value of not just the correct split, but also two incorrect splits. Thus, in this step the SVM is trained on four features, the Length of protein and the three interaction strengths, IS-Sum_2, IS-Sum_3 and IS-Sum_4, corresponding to the three splits, *k* = 2, 3, and 4, respectively.

### Step-3: Comparison with CATH Annotation

The number of domains, *k*, and the domain boundaries are compared with the CATH annotation. True prediction is reported if for proteins with correctly predicted number of domains, the fraction of correctly predicted residues is ≥ 75% compared to CATH annotation (Holland *et al* 2006). The algorithm also works if the user has prior information about the number of domains. In this case the steps – I and II and skipped and k-means algorithm is run on the given protein structure for user input *k*, the number of domains. The program outputs the domain boundaries.

### Implementation Details

We have developed a standalone program for the identification of structural domains in proteins by implementing the proposed algorithm, NML-DIP. Python scripts were written to parse the PDB files, construct protein contact network (PCN), compute feature vectors and apply SVM and k-means using Scikit-learn module (0.22.1). The algorithm works in two modes: the user may provide a file in PDB format (with or without the information about the chains) and the domain annotation (number of domains and domain boundaries) will be provided for all chains (or the specified chain). If the user is interested in identifying only the domain boundaries, then the user may provide a PDB file with chain ID and number of domains, k, as input. The algorithm is computationally very efficient and, on an Intel(R) Core (TM) i7-8565U CPU@1.80GHz system with 8 GB RAM, the algorithm took ∼ 2 - 3 seconds to execute and identify domains in a protein of size ∼ 450 residues.

## Results

### Dataset Construction

A brief description of training and test datasets used in this analysis is given below.

#### Training dataset

A dataset of 3000 proteins comprising 1500 chains each of single and multi-domain proteins is considered for training the first SVM. Further, this set of 1500 multi-domain proteins consists of 500 chains each of type 2-, 3- and 4-domain proteins and is used for training the second SVM. While constructing the dataset, care has been taken to include at least one representative domain defined by unique C, A and T categories in CATH (Orengo *et al* 1997), where C represents the secondary structure class of the domain, A the Architecture and T the 3-dimensional Topology. Also, it was ascertained that for the selected protein chains, the number of assigned domains agreed between CATH and SCOP.

#### Test datasets

Performance of the proposed integrated machine learning approach is evaluated on the following four test datasets, summarized in Table I.

**Table I.** The four test datasets used for performance evaluation are summarized.

| Test Datasets | 1-domain | 2-domain | 3-domain | 4-domain | Total |
| --- | --- | --- | --- | --- | --- |
| Benchmark_2 | 54 | 69 | 25 | 4 | 152 |
| Benchmark_3 | 55 | 53 | 21 | 3 | 132 |
| ASTRAL SCOP 30 | 4048 | 1599 | 489 | 154 | 6290 |
| NR_Dataset | 761 | 331 | 139 | 47 | 1278 |

#### Benchmark_2 and Benchmark_3 datasets

(Holland *et al* 2006): Benchmark_2 dataset consists of proteins chains for which domain assignment in CATH (Orengo *et al* 1997) and SCOP (Fox *et al* 2014) databases are in agreement and each protein chain is a representative of unique topology group in CATH (Orengo *et al* 1997). Benchmark_3 is a more stringent version of Benchmark_2 dataset, created by removing chains for which individual expert methods disagree with each other on the exact position of domain boundaries. Since only half of Benchmark_2 and Benchmark_3 datasets are made publicly available by the authors, both these datasets are small in size (152 and 132 proteins respectively), with very few 3-and 4-domain proteins.

#### ASTRAL SCOP 30 dataset

(Fox *et al* 2014): It is a non-redundant dataset at sequence level and consists of proteins having sequence similarity < 30%. Though it is the largest dataset considered (with 6290 protein chains), it is not truly non-redundant at the topological level.

#### Non-redundant dataset (NR_dataset)

To address the issues of class imbalance and redundancy in fair assessment of our algorithm, we constructed a non-redundant dataset (NR_Dataset), similar to the training dataset, in accordance with the approach proposed by Holland *et al* (2006). For its construction, protein chains for which the domain assignment agreed between CATH and SCOP were selected, resulting in 88,986 chains. These chains were grouped into topology classes represented by unique Class, Architecture and Topology in CATH, resulting in a total of 1313 topology groups. From each topology group, a *k* domain protein was randomly picked, where *k* = 1 - 4. For single domain proteins, selection is simple and straightforward. For multi-domain proteins, selection was done such that at least one of the domains of the protein is a unique representative of each topology group. A step-by-step evaluation of our algorithm is provided on this non-redundant NR_dataset. The comparison of the performance of various tools is carried on Benchmark_2, Benchmark_3 and Astral SCOP 30 datasets.

#### Single *vs* Multi-domain Classification

The first SVM is trained for distinguishing between single and multi-domain proteins. The results of SVM-I on NR_dataset is summarized in Table II. The overall prediction accuracy of ∼ 84% is observed in single *vs* multi-domain classification. We observe that the length of protein is a very crucial parameter and majority of incorrectly classified single domain proteins are much larger in size compared to the average single domain length (∼ 154). In Figure 2 representative examples of incorrectly assigned single and multi-domain proteins are depicted. As seen in Figure 2 (A), the misclassified single domain protein 1XIM (A) is of length 394, much larger than the single-domain average. Similarly, we observe that 4 (Benchmark_2), 11 (Benchmark_3), 183 (ASTRAL SCOP 30) and 82 (NR_Dataset) single-domain proteins that were wrongly classified as multi-domain proteins have significantly larger length compared to single-domain average.

**Table II:** Assessment of various steps in the proposed algorithm on the NR_Dataset is shown. Here the numbers in bold correspond to the overall percentage of correctly classified proteins, C and NC that of contiguous and non-contiguous multi-domain proteins, respectively.

| Protein | No. of Proteins | SVM-1 | Multi-Domain Proteins |  | SVM-2 | Prediction of Domain Boundaries |
| --- | --- | --- | --- | --- | --- | --- |
|  |  |  | No. of Proteins | No. of Domains |  |  |
| Single | 761 | 92.77 | - | - | - | - |
| Multi | 517 | 78.92<br>(C: 76.4, NC: 82.2) | 331 | 2 | 96.65<br>(C: 96.5, NC: 96.8) | 87.01<br>(C: 87.05, NC: 87) |
|  |  |  | 139 | 3 | 63.11<br>(C: 62.9, NC: 63.3) | 76.62<br>(C: 82, NC: 75) |
|  |  |  | 47 | 4 | 80.85<br>(C: 88.2, NC: 76.7) | 86.84<br>(C: 86.7, NC: 87) |
| Total | 1278 | 87.17 | 517 | - | 84.80<br>(C: 86.5, NC: 82.7) | 84.68<br>(C: 86, NC: 83) |

**Fig. 2:**
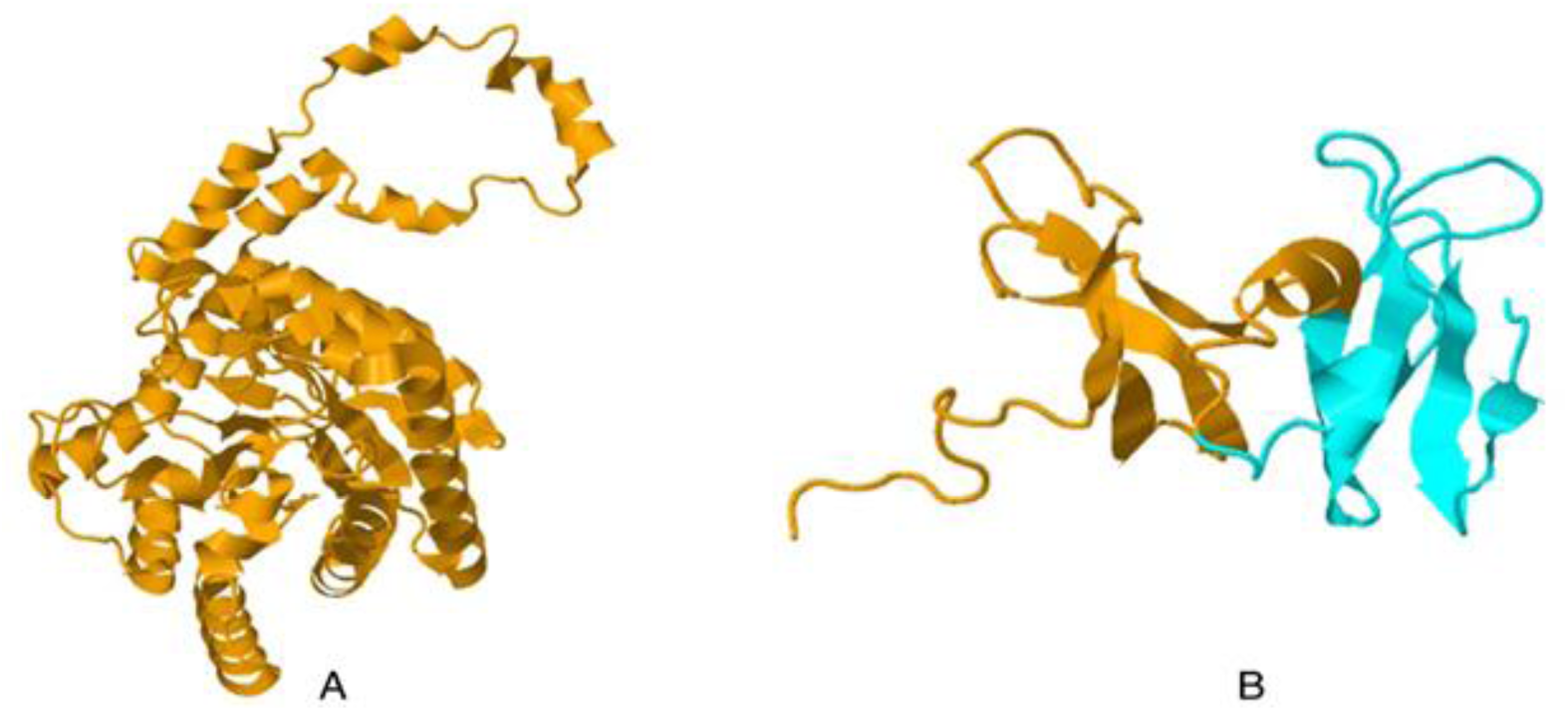
(A) Protein 1XIM (chain A) is a single domain protein of length 394, much larger than a typical single domain protein. Also, it may be noted that the protein does not fold into a very compact 3D structure resulting in lower values of Density and Interaction Strength, 0.039 and 3.973 respectively, compared to the respective average values for single-domain proteins, ∼ 0.047 and ∼ 11.63, respectively. (B) Protein 1YUA (chain A) is a two-domain protein, Domain I:1-64 and Domain II: 65-122. It’s a very small protein of length 122, much smaller than the average length of multi-domain proteins, ∼ 434.

Apart from length, compactness is also observed to be an important feature in domain identification and is captured in our SVM through the graph properties, density and interaction strength. A number of single domain proteins wrongly classified in the four test datasets have lower Interaction Strength and/or lower density than the average value for single-domains. For e.g., apart from being a large protein, 1XIM (A) has lower density (= 0.039) and interaction strength (=3.973) values compared to the average for single-domain proteins, ∼ 0.047 and ∼ 11.63, respectively. Similarly, it is observed that most wrongly predicted multi-domain proteins are smaller in size compared to the average (∼ 434). The For example, protein 1YUA (chain B) shown in Figure 2 (B) is of length 122, much smaller than the multi-domain average. We observe 8 (Benchmark_2), 6 (Benchmark_3), 108 (ASTRAL SCOP 30) and 34 (NR_Dataset) misclassified multi-domain proteins that have lengths smaller than the average.

#### Identifying Number of Domains

In the second step of the algorithm, a second SVM is used in *de novo* detection of number of domains in multi-domain proteins. It is trained on four features: the Length of protein (L), and three interaction strengths, IS-Sum_2, IS-Sum_3 and IS-Sum_4, corresponding to the sum of inter-cluster interactions on splitting the protein in two, three and four clusters using k-means algorithm. It may be noted from Table II that prediction accuracy of the second SVM in correctly identifying the number of domains on NR_Dataset is ∼ 82%, comparable to the first SVM. It is worth noting that the performance is equally good for non-contiguous as for contiguous proteins, while many algorithms fail to detect non-contiguous domains. The performance of our algorithm is good in the detection of 2-domain (∼93%) and 4-domain (∼83%) proteins. However, the prediction accuracy dropped to ∼60% for 3-domain proteins with majority of them incorrectly labeled as 2-domain proteins. On closely examining these proteins, we observed two features of 3-domains coupled with limitations of k-means algorithm in forming clusters of similar sizes that resulted in poor prediction accuracy on 3-domain proteins. Below we discuss these features with few representative examples of 3-domain proteins.

#### Size of a domain ≥ sum of remaining domains

For a number of 3-domain proteins incorrectly predicted as 2-domain, the size of one of the domains is observed to be comparable or larger than the combined size of remaining two domains. In Figure 3 (IA) is shown the 3-domain protein 1BHG (A) with CATH annotation as Domain I: 22-224 (size: 202), Domain II: 225-328 (size: 103) and Domain III: 329-631 (size: 302). It may be noted that the size of Domain III (302) is comparable to sum of the size of domains I and II (= 305). Since k-means algorithm tends to generate clusters of similar sizes while simultaneously maximizing the distance between centroids of each cluster, the IS-Sum_2 value of 1BHG is similar to that of 2-domain proteins and the algorithm classifies it as a 2-domain protein, merging the two smaller domains as depicted by ‘red’ colour in Figure 3 (IB). Similarly, 3-domain proteins 1RB5 (A), 2RE3 (A), 1PIX (A), shown in Figure 3 II, III and IV respectively, also exhibit a similar pattern with size of one of the domains much larger than the other two domains.

**Fig. 3:**
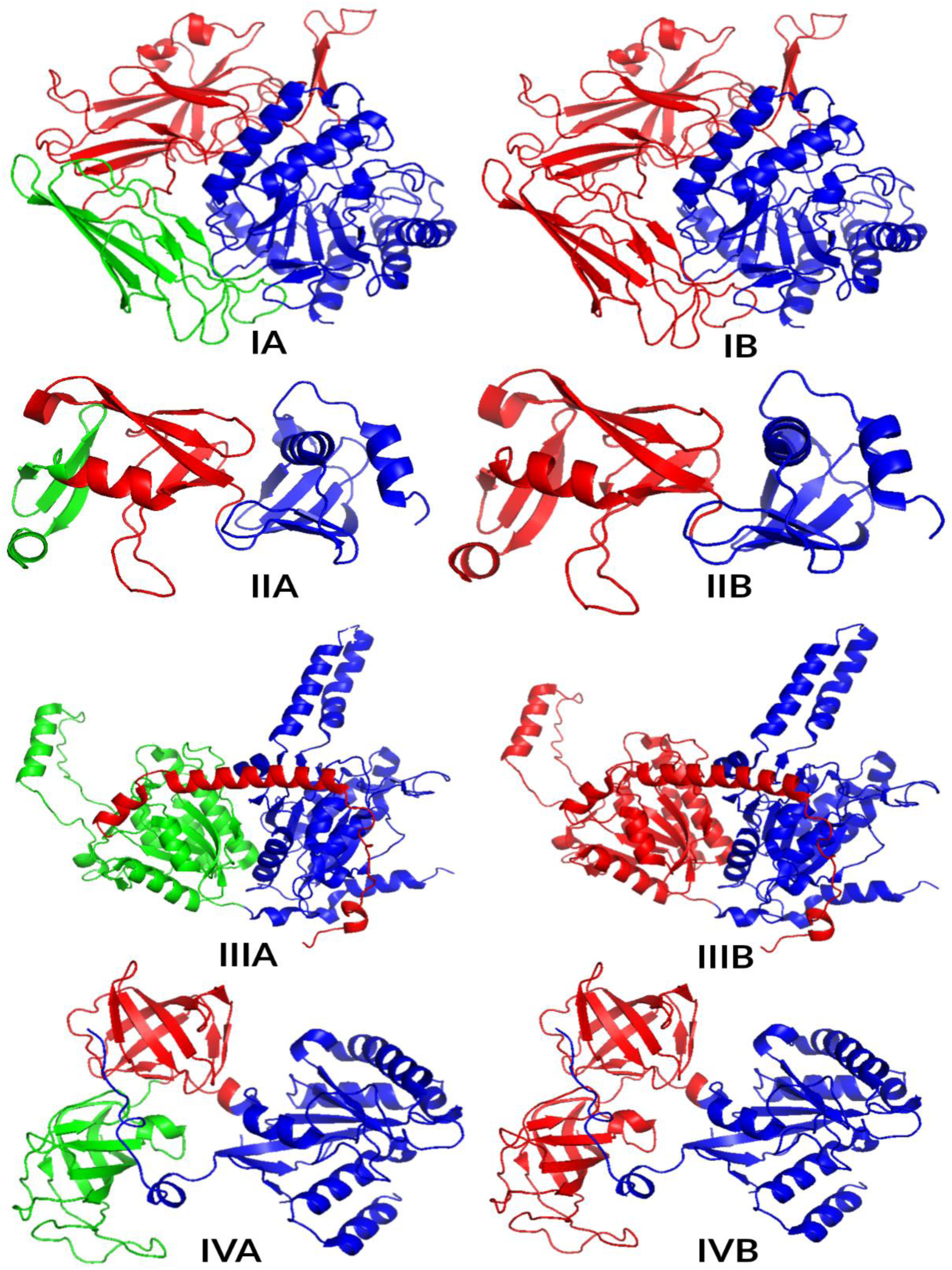
Representative examples of 3-domain proteins in which size of one domain is larger or comparable to the combined size of remaining two domains is shown. I: 1RB5 (A), II: 2RE3 (A), III: 1PIX (A) and IV: 1BHG (A). In the left column (labeled ‘A’) CATH annotation is depicted with three domains shown in three different colours, ‘red’, ‘green’ and ‘blue’. In the right column (labeled ‘B’) the prediction from our algorithm is depicted with two smaller domains merged and represented in ‘red’ colour.

#### One of the domains is spatially distant from other two

In Figure 4 representative examples of 3-domain proteins are shown, wherein one of the domains, depicted in ‘blue’ is spatially distant from the remaining two domains shown in ‘red’ and ‘green’ colour. Being in close proximity, the number of interactions between ‘red’ and ‘green’ domains is much higher compared to that between ‘red’ and ‘blue’ domains, or ‘green’ and ‘blue’ domains. This results in IS-Sum_2 values comparable to that of average 2-domain proteins and the SVM ends up classifying them as 2-domain proteins.

**Fig. 4:**
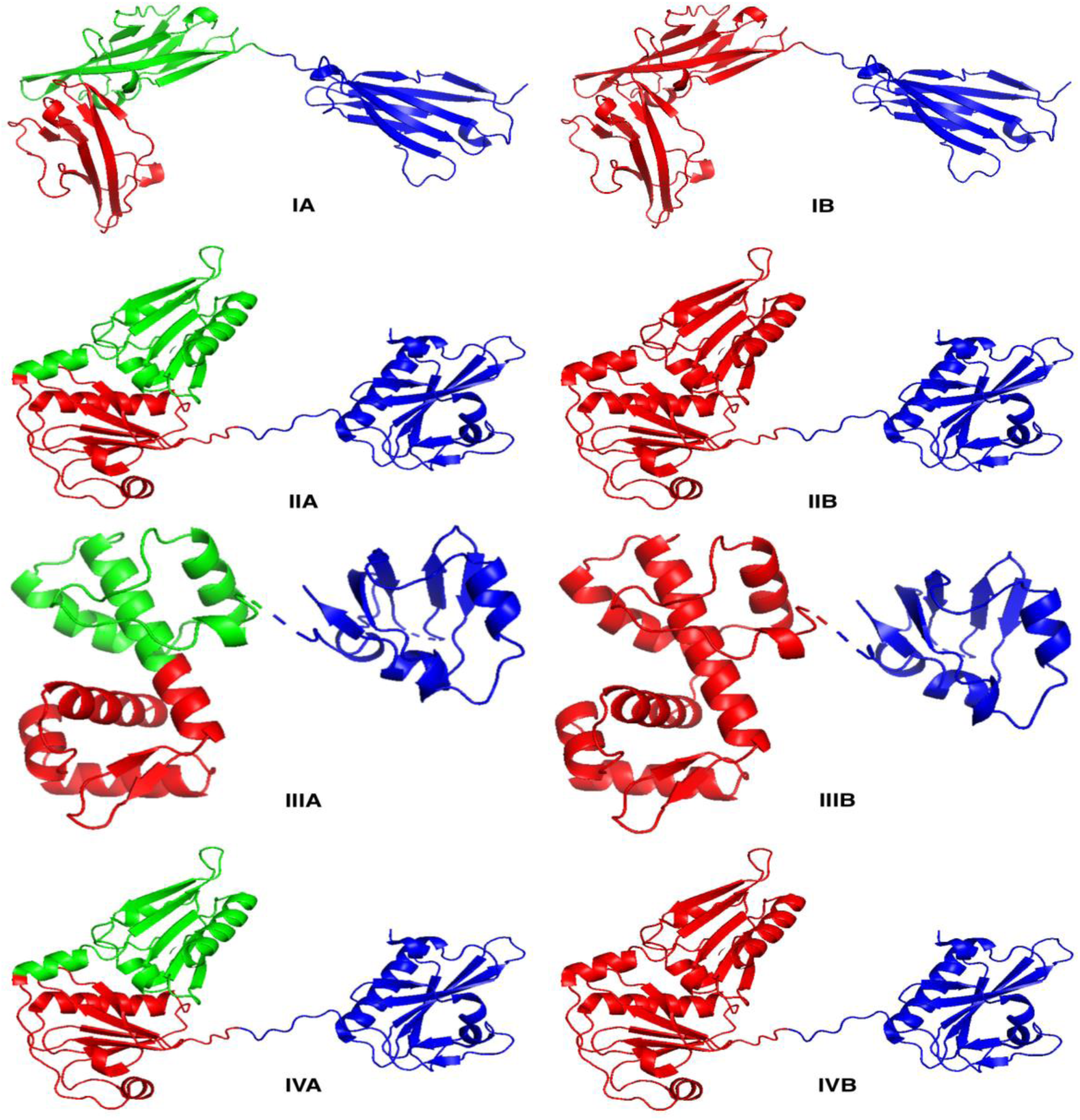
Representative examples of 3-domain proteins with one domain being spatially distant from other two as shown in I: 1IRA (chain Y), II: 3FFK (chain A), III: 1BI3 (chain A), IV: 3C18 (chain A). In the left column (labeled ‘A’) CATH annotation of 3-domain protein is depicted and in the right column (labeled ‘B’) the prediction from our algorithm is shown as 2-domain protein.

#### Comparison of Domain Boundaries with CATH

The proteins for which the number of domains are correctly identified in the previous step, domain boundaries are extracted for the correct *k*-split and true prediction is reported if the fraction of correctly predicted residues is ≥ 75% compared to CATH annotation. On using k-means, sometimes clusters with very short fragments (< 20 residues) are observed. Suppose, for example, the k-means output for a 2-domain protein is: Domain 1: 1-70, 81-100, Domain 2: 71-80, 101-180. In such a case the fragment 71-80 is merged with domain 1, resulting in two domains as Domain 1: 1-100, Domain 2: 101-180. This post-processing is performed only for short segments < 20 residues, since even for non-contiguous domain proteins, fragments shorter than 20 residues are not observed. The results are summarized in the last column of Table II. The k-means clustering algorithm performed quite well with an overall prediction accuracy of ∼ 90% on the NR_Dataset. Thus, we see that a simple unsupervised algorithm such as k-means is able to capture very well the structural features of domains.

## Comparison of Results with Other Domain Identification Methods

We compared the prediction accuracy of our integrated approach with four state-of-the-art structure-based domain identification tools, namely, CA Algorithm (Feldman 2012), DDomain (Zhou *et al* 2007), DomainParser2 (Guo *et al* 2003) and PDP (Alexandrov and Shindyalov 2003) on three datasets: Benchmark_2, Benchmark_3 and ASTRAL SCOP 30. The α-carbon based algorithm (CA) uses a very simple approach to identify structural domains and works well on individual chains or protein complexes. It clusters buried alpha carbons using average-linkage clustering to produce and cut a dendogram to identify structural domains. The DDomain algorithm identifies domains by dividing a protein chain using a normalized contact-based domain-domain interaction profile under the assumption that inter-domain interactions are the weakest under the correct assignment. DomainParser2 uses a top-down graph theoretical approach for domain decomposition, formulating it as a network flow problem and solves it using Ford–Fulkerson algorithm. Its performance is best when the number of domains is provided. Protein Domain Parser (PDP) uses a similar approach to identify domain by hierarchical decomposition of a protein into smaller fragments based on the idea that inter-domain contacts are much more than intra-domain contacts.

Here a prediction is considered to be true if the number of domains is correctly identified and the predicted domain boundaries have an overlap of ≥ 75% with CATH annotation. The overall accuracy corresponds to the average accuracy in detecting 1, 2, 3 and 4-domain proteins correctly. It may be noted from Fig. 5 that the performance of all the algorithms is comparable with small differences across the three datasets. While CA algorithm and PDP perform better on benchmark_2 and benchmark_3 datasets, our algorithm and DDomain perform better compared to other on ASTRAL SCOP 30 dataset, which is much larger than the two benchmark datasets. Our algorithm’s performance is observed to be comparable with other methods in identification of single, 2- and 4-domain proteins while the performance dips for 3-domain proteins. This may possibly be due to the features of the 3-domain proteins discussed above.

**Fig. 5:**
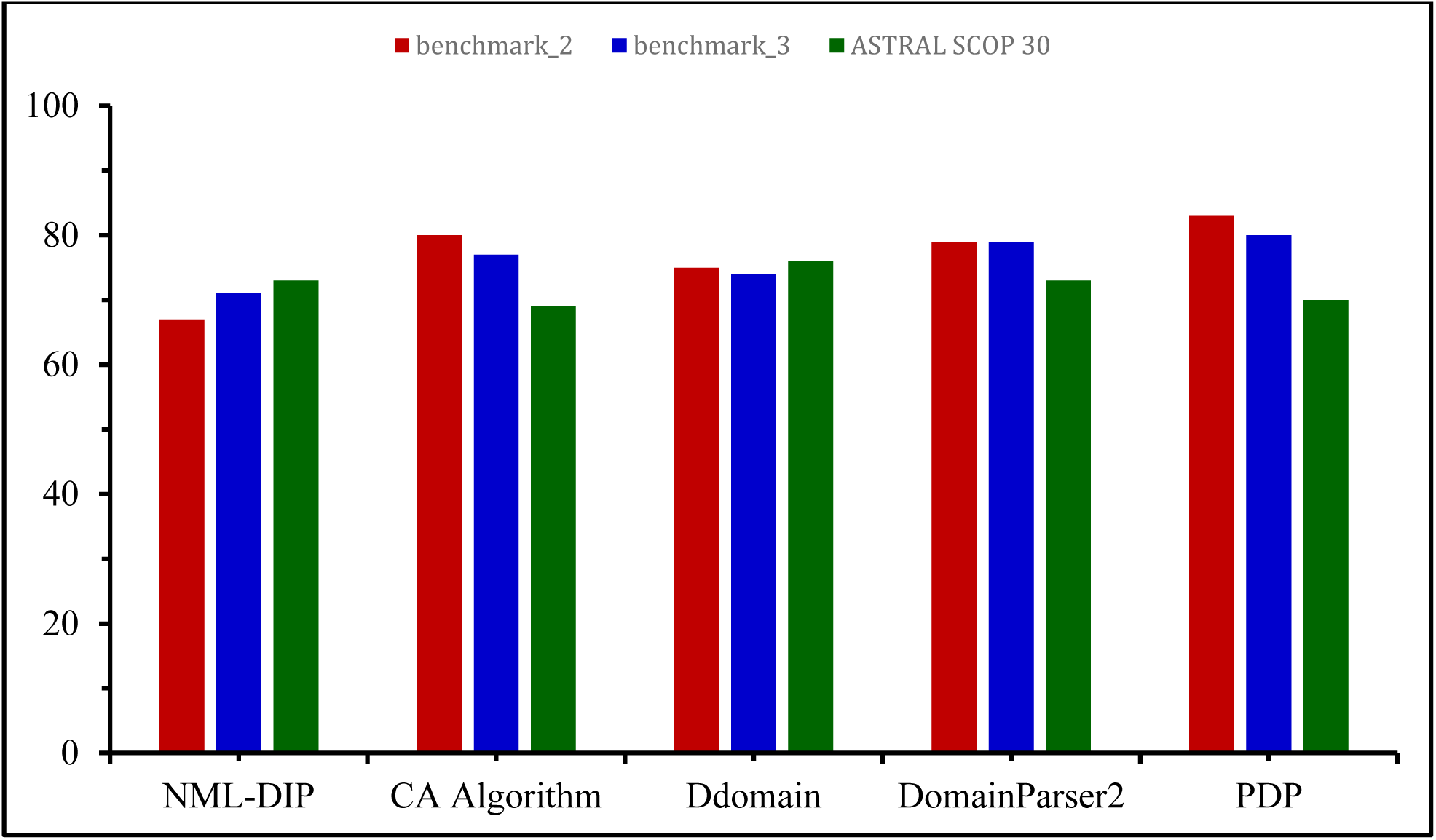
Comparing overall domain prediction accuracy of four state-of-the-art algorithms with NML-DIP on three datasets, namely, benchmark_2, benchmark_3 and ASTRAL SCOP 30. The results for the four state-of-the-art algorithms is reproduced from CA Algorithm (Feldman 2012).

We also present the overall performance of NML-DIP on the non-redundant NR_Dataset in Table III. It may be noted that the overall accuracy in detecting single domains in NR_Dataset is comparable with those on benchmark_2 and benchmark_3 datasets (73-74%). The overall prediction accuracy of 3-domain proteins is also improved on this dataset. This clearly indicates how the test dataset can affect the results and a large non-redundant dataset is clearly required for assessing the performance of the algorithm.

**Table III:** Performance of NML-DIP on NR_Dataset.

|  | 1-domain | 2-domain | 3-domain | 4-domain | Overall |
| --- | --- | --- | --- | --- | --- |
| **NML-DIP** | 92.77% | 60.73% | 42.45% | 68.09% | 78.09% |

## Conclusion

It may be noted that the domain identification problem is very similar to community detection in a social network in the sense that the number of domains or communities is not known a priori and also both exhibit large number of connections within a community/domain than between communities/domains), allowing us to borrow techniques from social network theory to address biological problems. Further, since numerous definitions have been proposed to define a domain, an approach not dependent on the domain knowledge is desirable. With this observation we proposed here a combination of graph theory (for feature selection) and machine learning approach for domain identification. We show that using graph properties such as length, density and interaction strength as feature vectors in SVM algorithm, we are able to distinguish between single and multi-domain proteins, as well as multi-domain protein into k-domains. Here, Interaction Strength is a novel feature vector which captures both inter- and intra-domain interactions based on spatial proximity. The prediction accuracy of our algorithm on ASTRAL SCOP 30 dataset is comparable with other tools. This suggests that a combination approach can really help in improving the sensitivity of domain detection algorithms.

